# Thermal transients excite neurons through universal intramembrane mechano-electrical effects

**DOI:** 10.1101/111724

**Authors:** Michael Plaksin, Eitan Kimmel, Shy Shoham

## Abstract

Modern advances in neurotechnology rely on effectively harnessing physical tools and insights towards remote neural control, thereby creating major new scientific and therapeutic opportunities. Specifically, rapid temperature pulses were shown to increase membrane capacitance, causing capacitive currents that explain neural excitation, but the underlying biophysics is not well understood. Here, we show that an intramembrane thermal-mechanical effect wherein the phospholipid bilayer undergoes axial narrowing and lateral expansion accurately predicts a potentially universal thermal capacitance increase rate of ~0.3%/°C. This capacitance increase and concurrent changes in the surface charge related fields lead to predictable exciting ionic displacement currents. The new theory’s predictions provide an excellent agreement with multiple experimental results and indirect estimates of latent biophysical quantities. Our results further highlight the role of electro-mechanics in neural excitation; they may also help illuminate sub-threshold and novel physical cellular effects, and could potentially lead to advanced new methods for neural control.

---

Optical neurostimulation modalities have gained considerable attention during the past decade as methods for precision perturbation or control of neural activity, primarily as a result of the co-emergence of optogenetics^1^ and of direct infrared neural stimulation (INS)^2^. Both approaches also offer the long-term prospect of remotely affecting aberrant localized neural circuits that underlie many neurological diseases. A multitude of INS-related studies explored the ability of short wave infrared (IR) pulses to stimulate neural structures including peripheral^3, 4^ and cranial nerves^5–10^, retinal and cortical neurons^10–12^, as well as cardiomyocytes^13, 14^ It is stipulated that the INS phenomenon is mediated by temperature transients induced by IR-absorption^15–17^; such transients can alternatively be induced using other forms of photo-absorption^18–20^, or potentially by any other physical form of thermal neurostimulation that can be driven rapidly enough^21, 22^. Shapiro *et al.* (2012)^16^ showed that these rapid temperature variations are directly accompanied by changes in the cell membrane’s capacitance and resulting displacement currents which are unrelated to specific ionic channels. Shapiro *et al.*^16^ also developed a theoretical model where the temperature elevation was seen to give rise to membrane capacitance increase at the membrane’s boundary regions (see also Liu *et al.* 2014^23^ and Rabbit *et al.,* 2016^24^). In these theoretical models, however, the projected capacitance *increase* in the membrane boundary region as the temperature increases is seemingly paradoxical; from energetic considerations for example, thermal energy input will be offset by correspondingly higher electrical energy and absolute potentials, which corresponds to a capacitance *decrease,* and this decrease is further compounded by a temperature-related decrease in the dielectric constant of water. Indeed, a reanalysis of their model (see below), quantitatively predicts such a net capacitance decrease, which is contrary to the experimental measurements, showing that the seemingly complete theoretical picture resulted from a mathematical convention error.

To address this major apparent gap in the theoretical basis for thermal excitation, we deconstruct and analyze here alternative revised biophysical models (tentative and detailed), where the membrane’s physical dimensions themselves also vary in response to the temperature changes, to accurately reflect direct experimental findings^25–27^. The new models calculate the effect of temperatures change on the membrane electrical parameters directly and explicitly (rather than implicitly), and are found to both qualitatively and quantitatively predict empirical findings on thermal membrane capacitance increases^16, 19, 20, 28^, unraveling the two underlying sources for thermal membrane currents, and making it possible to indirectly estimate from neural stimulation results a significant new quantity, the membrane surface charge difference.

## Results

### Dimensional changes are crucial for explaining capacitive thermal response

Capacitive thermal changes were first measured by Shapiro *et al.*^16^ and subsequently by others in artificial membranes^19^, HEK cells^20^ and cardiac myocytes^28^. Interestingly, placing these disparate measurements on a uniform capacitance-rate scale, shows that they all approximately share a universal rate of ~0.3 %/°C (mean: 0.29±0.01 %/° C), suggesting a potentially universal basis (Fig. 1a).

**Figure 1.**
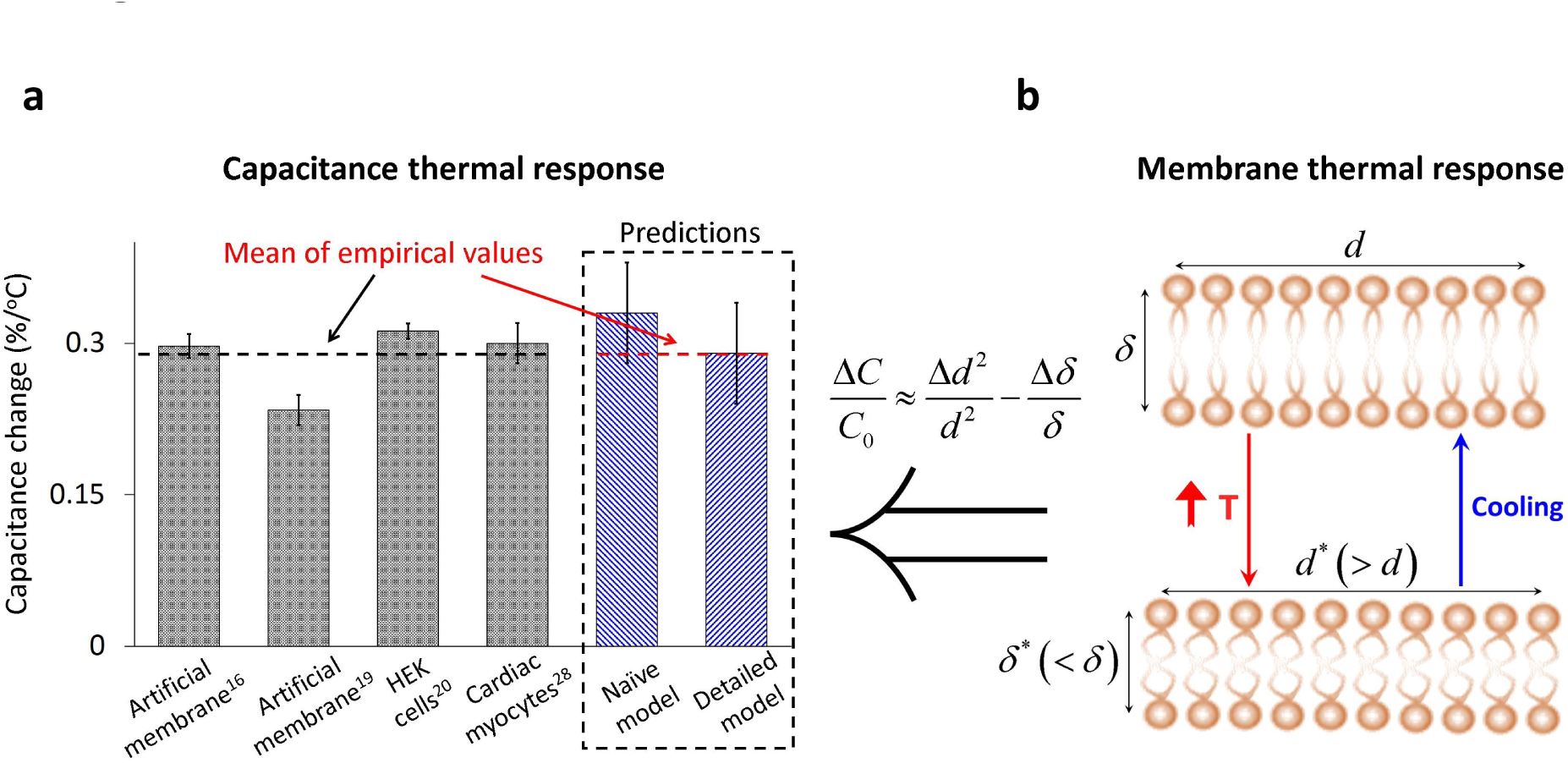
Universal membrane capacitance thermal response rate, explained by membrane dimensional changes. (**a**) Membrane capacitance change rates measured in different studies and preparations (gray bars) vs. predictions from the thermal dimensional response of POPC membrane bilayers (Naive plate capacitor model), and from a detailed biophysical model accounting for spatial charge distribution. (**b**) Schematized membrane thinning and area expansion under temperature elevation. This observed process^25–27^ is thought to result from an increase in the phospholipid molecules’ fatty acid chains transgauche rotational isomerization, which shortens the tails’ effective length and increases the area per phospholipid molecule. Biophysically, these two phenomena contribute to a predictable increase in both the membrane hydrophobic core’s and total capacitance. Error bars for the direct capacitance measurements are ±s.e.m, and for the model predictions are ± chi-squares distribution-related uncertainty^26^.

We next examined how these changes compare to the membrane’s capacitance thermal rate-of-change expected purely from temperature-induced dimensional changes, which were recently experimentally estimated using X-ray and neutron small-angle scattering measurements^25–27^. Using a naive plate capacitor assumption where 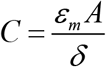, the 0.11±0.03 %/°C reduction in the phospholipid membrane thickness and 0.22±0.03 %/°C increase in the area per phospholipid molecule (Fig. 1b) contribute to a linear increase of 0.33±0.05 %/°C (relative to the corresponding parameters in 20°C, assuming that the membrane’s dielectric constant is temperature independent^29^), putatively explaining the observed capacitance rates (Fig. 1a).

### Detailed biophysical model

To further understand the impact of this mechanism on membrane currents and its potential for settling the theoretical conceptual gap, we subsequently studied the effect of temperature changes and transients on detailed Gouy-Chapman-Stern^30–34^ (GCS) multi-compartment realistic biophysical model of the phospholipid membrane electrochemistry. The various model parameters were taken “as is” without re-tuning or post-hoc adjustments, and are based on known or previously measured physical and biophysical quantities, wherever attainable (summarized in Supplementary Table 1 with the respective sources). In the model, the membrane geometry and physical properties are represented by five regions with different characteristics (Fig. 2a): two bulk regions for each of the membrane’s sides (intracellular and extracellular), where the ions’ concentrations are spatially Boltzmann-distributed, obtained when diffusion and electrical forces on the different ions reach equilibrium; two Stern layers with thicknesses determined by the phospholipid polar head size and the average effective radius of the ions that are the closest to the membrane and a central hydrophobic region, occupied by the tails of the phospholipid molecules^16, 30–34^. The ions in each of the bulk regions comprise of the mobile and fixed extra-membranal charges (intracellular: *Q* and –*σ*_*i*_; extracellular: *–Q* and *–σ_o_*); the fixed charges have the same absolute value, but opposite signs as the membrane leaflets’ surface charges to which they are electrostatically attracted (Fig. 2a). The relationships between the mobile, fixed charges and the potentials that fall on each membrane region are determined by electrostatic Boltzmann equations in the bulk, while in the hydrophobic core and Stern layers these relations are determined through the assumption of zero density of free charges^16, 33, 34^ (see Methods). Interestingly, approximately linear relations between the mobile charge and the potentials are found in all five membrane regions (Supplementary Fig. 1), allowing the membrane to be represented as a lumped circuit of 5 temperature-dependent capacitors with surface charge-related sources in series (alternatively, as a single equivalent temperature dependent membrane capacitor and surface charge-related source, see Fig. 2b and Methods section).

**Figure 2.**
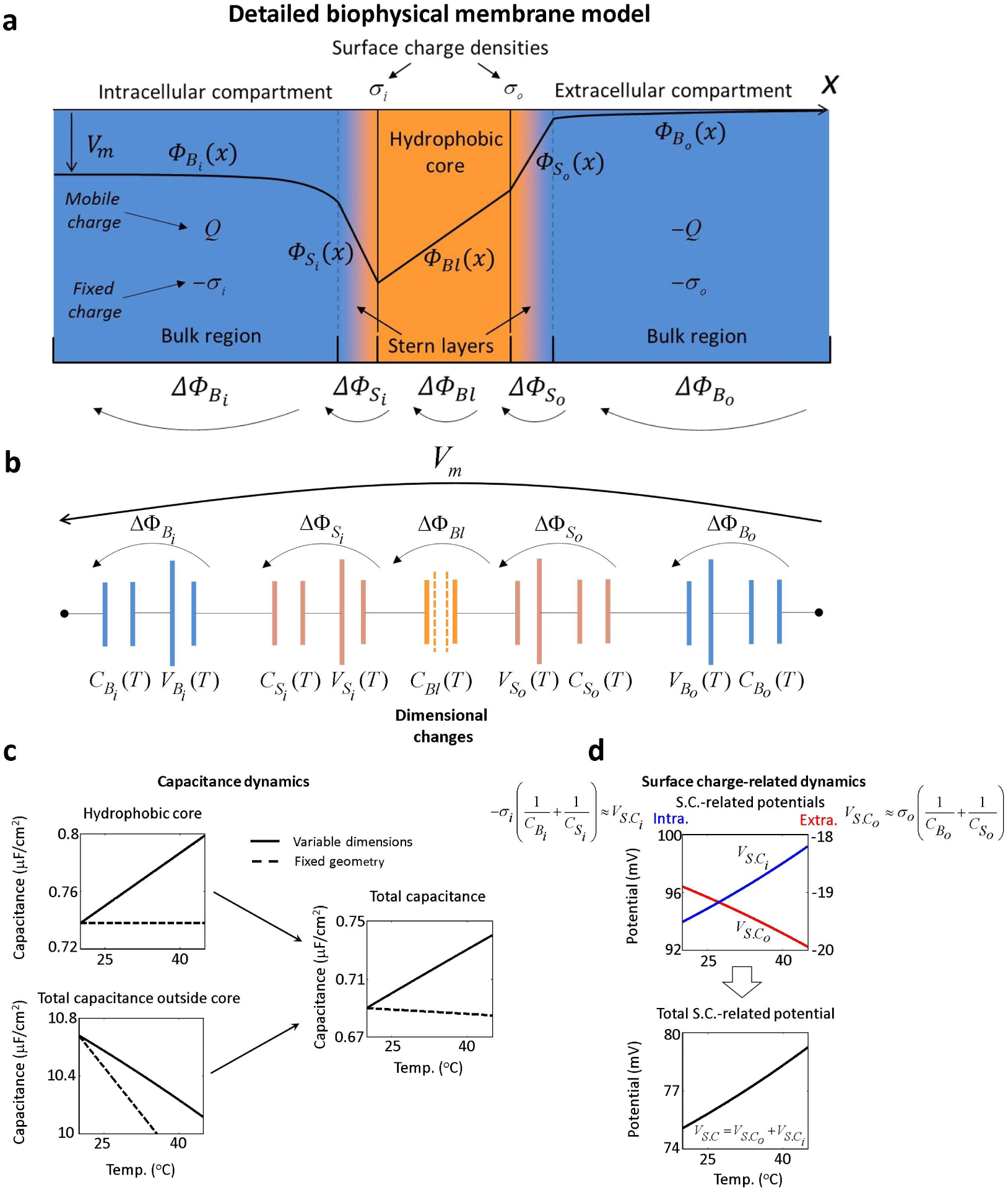
Detailed (Gouy-Chapman-Stern-based) membrane biophysical model. (**a**) Membrane sub-regions: *hydrophobic core,* containing the phospholipid molecules’ tails. The outer parts of this region contain intracellular (σ_i_) and extracellular (σ_o_) surface charges. *Intracellular and extracellular bulk regions* containing mobile charges (Q and −Q) and the surface charges’ fixed counterions (−σ_i_ and −σ_o_). *Two Stern layers* comprised of the phospholipid polar heads and the closest ions to the membrane. *ΔΔΦ_j_* is the potential drop on each sub-region (Total: *V_m_=Σ Δ Φ_j_).* (**b**) Membrane series circuit where capacitors and sources approximate each sub-region (*Δ Φ_j_ = Q/C_j_ + V_j_*)*. V_j_*≈ −*σ_i_;* C_*j*_ for *j=B_i_* or *S_i_* and *V_j_* ≈ σ_*o*_/C_*j*_ for *j=B_o_* or *S_o_* is the potential resulting from respective surface charges and both *C_j_* and *V_j_* are temperature dependent. Overall, *Q=C_m_*(*V_m_-V_s.c_*), where *V_s.c_*. is the surface charge-related potential. (**c**) Predicted thermal changes in partial and total membrane capacitance under alternative models: fixed geometry model predicts a net reduction while the varying-dimension model correctly predicts an increase (see also Fig. 1a). (**d**) Predicted thermal response of intracellular, extracellular (upper panel) and total (bottom panel) surface charge-related potentials.

We first examined the fundamental response to temperature changes of the new membrane biophysical model and of an equivalent fixed-geometry model^16^. The fixed-geometry model predicts a paradoxical net *reduction* with temperature of the total membrane capacitance (−0.03 %/°C) dictated by a reduction of the capacitance outside the membrane core (−0.4 %/°C, Fig. 2c, see Discussion and Methods for explanation of earlier misinterpretations^16, 23^). In contrast, in the new model, the membrane’s hydrophobic core dimensional variations (Fig. 1b) contribute to a linear increase of 0.29±0.05 %/°C in the total membrane capacitance (Fig. 1a and Fig. 2c, right panel), which is dominated by this relatively small capacitor, a value that accurately matches the mean experimental capacitance change rate (Fig. 1a). Importantly, although the Stern and bulk capacitances actually undergo a concurrent temperature-dependent *drop* (Fig. 2c) caused by the reduction in the surrounding dielectric constants and an elevation of the membrane’s thermal/electrical forces, this effect is responsible for another surface charge-related phenomenon: an increase in the absolute values of the potentials falling on the Stern and bulk regions, contributing to an overall elevation of the total surface charge-related potential *V_S_C_* (Fig. 2d).

### Experimental validation and inference

We next studied the model’s response to a temperature *transient* used extensively by Shapiro *et al.* in their artificial membranes’ experiments and model simulations (Fig. 3 a). The resulting membrane currents and potential changes were composed of similar relative contributions from the two underlying mechanisms related to capacitance and surface charge-related potential variations (Fig. 3b,c). Importantly, the membrane capacitance-related current is dependent on the clamped membrane potential, while the surface charge-related current is independent (Fig. 3b).

**Figure 3.**
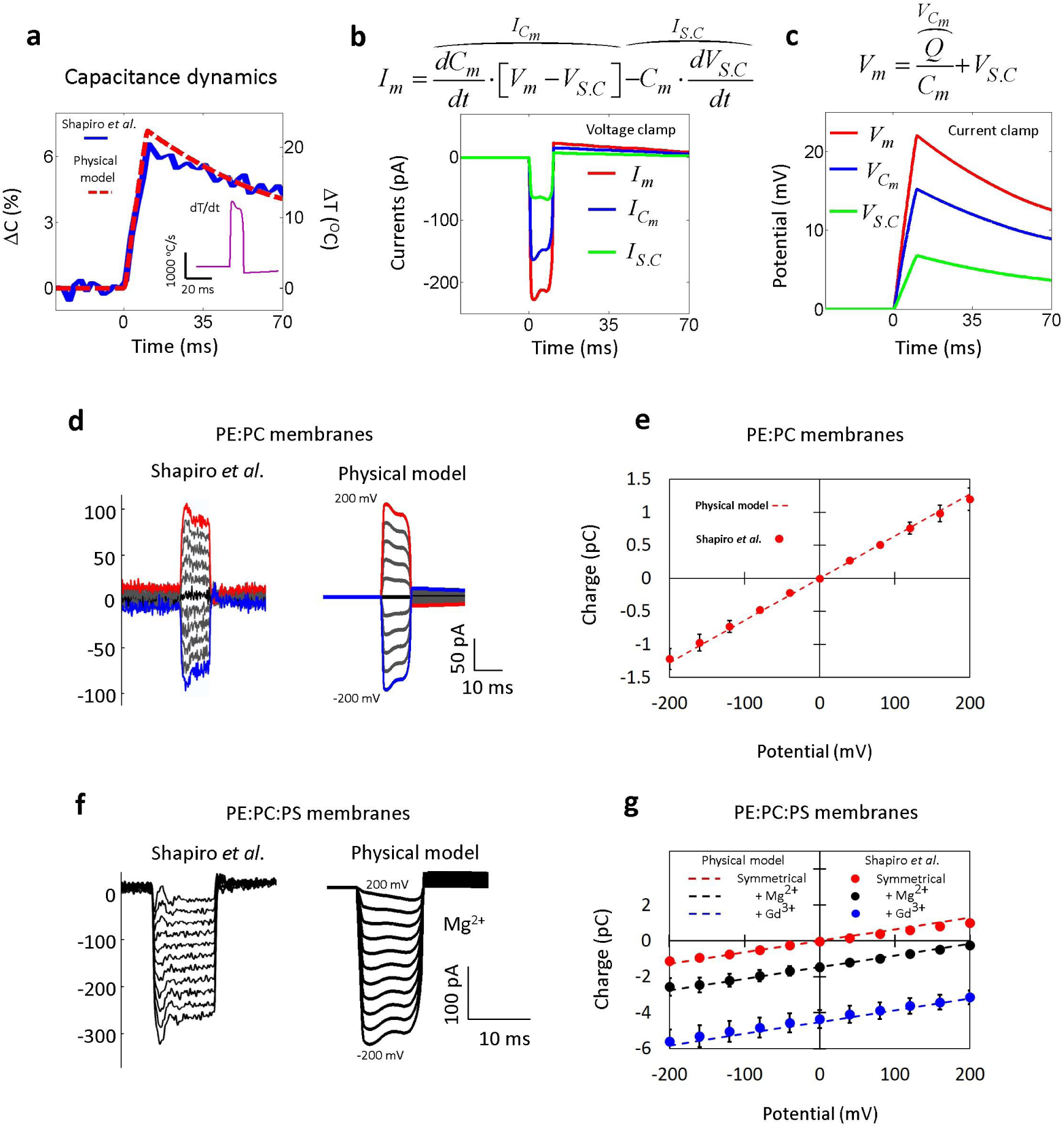
Empirical results vs. model-based predictions for artificial bilayer membrane INS experiments. **(a)** Predicted vs. measured capacitance dynamics for 10 ms pulses. The capacitance change is proportional to the temperature transient (inset - temperature time derivative). (b, c) Underlying membrane currents and potentials in response to the temperature transient in *(a).* The overall response is determined by the additive contribution of the membrane capacitance and surface charge-related potential thermal responses. **(d-g)** Predicted vs. measured thermal membrane currents *(d,f)* and induced charge *(e,g)* for PE:PC membranes and for PE:PC:PS membranes. Addition of 14 mM MgCl_2_ *(f,g)* or 1mM GdCl_3_ *(g)* to the “extracellular” compartment, shifts towards more depolarizing currents through membrane absorption of excess ions changing the extracellular surface charge density. The simulations use Shapiro *et al*.^16^ experimental parameters (Supplementary Table 1). Error bars are ±s.e.m.

Can these dynamical responses quantitatively predict the results of INS experiments in artificial bilayer membranes^16^? In simulations of PE:PC or PE:PC:PS membranes (Energy: 7.3 mJ; Duration: 10 msec, see Methods), the resulting INS temperature transient caused a membrane capacitance increase of about 0.3%/°C, in agreement with the respective experimental measurements^16^ (Fig. 3a). The same high level of agreement was also obtained in voltage clamp simulations for simulated membrane currents and temperature elevation phase-induced membrane charge (*Δ*Q) in PE:PC (Figs. 3d,e) and PE:PC:PS (Figs. 3f,g) membranes. For PE:PC:PS membranes where multivalent ions were added to the pseudo “extracellular compartment” (Mg^2+^ or Gd3+, as 14mM MgCl_2_ or 1mM GdCl_3_, see Methods for modeling detail), the membrane currents were shifted towards more depolarizing currents due to changes in surface charge difference between the extracellular and intracellular compartments. This surface charge difference elicited a depolarizing current (shown earlier in Fig. 3b), which was absent when the solutions on both sides of the membrane were symmetrical (Fig. 3f, g).

Our final analyses examined whether the new biophysical model can also predict and explain thermo-stimulation results in intact living cells. First, we simulated INS stimulation experiments performed in Xenopus laevis oocytes^16^ (see Supplementary Table 1 for simulation parameters), finding a high degree of agreement between the model predictions and the experimental measurements of both currents and potentials (Fig. 4a,b). Next, we performed a model-based re-analysis of our recent experiments where thermal excitation mediated by photo-absorption of CW laser pulses with varying durations was used to excite cells in rat cortical cultures^18^. Farah. *et al*.^18^ observed stimulation thresholds are well-predicted if putative depolarizing displacement currents proportional to the temperature’s time derivative 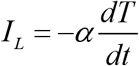 (α=2.53*10^−5^ Coulomb/°C/m^2^, Fig. 4d, based on ref. 18 Fig. 5a) are inserted into model layer V cortical neurons. Re-examining in terms of the new biophysical model, α is almost entirely dependent on the temperature derivative of the membrane capacitance multiplied by surface charge-related potential, allowing to estimate the surface charge difference, Δσ= σ_i_−σ_ο_, as −0.107 C/m^2^ plus maximal deviation ≈ 5% (see Methods for detail). This result is consistent with the surface charge density range of 0.002−0.37 C/m^2^ that was measured in neural cells^35^.

**Figure 4.**
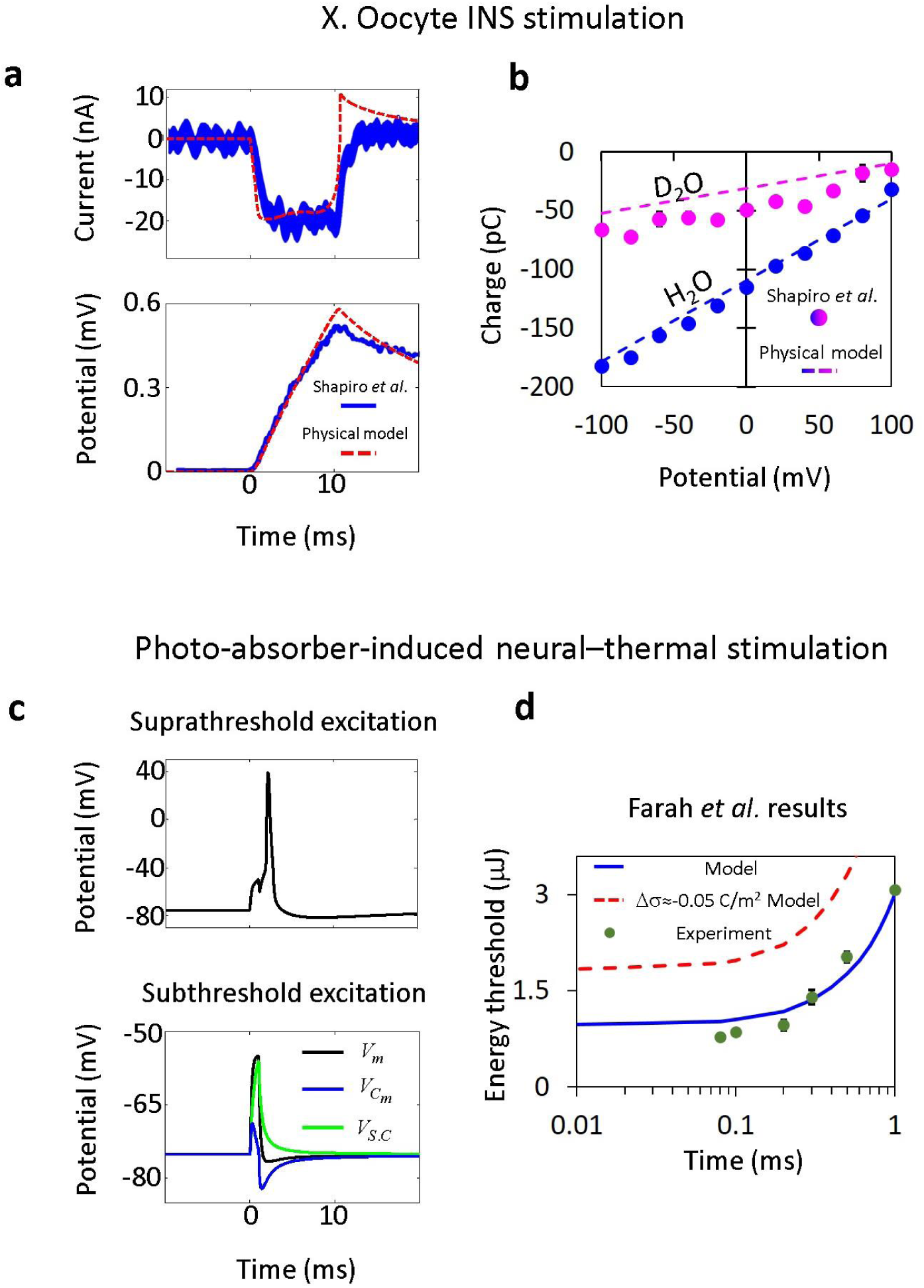
Empirical cellular neuro-stimulation results vs. model-based predictions. (**a,b**) Predicted vs. measured membrane currents, potentials and induced charge, for 10ms 7.3 mJ INS stimulation of Xenopus oocyte in H_2_O and D_2_O-based extracellular media; illumination of 5% of the surface area assumed^16^. (**c**) Predicted somatic membrane potentials in a cortical pyramidal neuron stimulated by threshold and 90% sub-threshold simulated 1 ms PAINTS transients. The membrane potential (*V_m_*) thermal response is dictated by surface charge and capacitance-related potentials (*V_S.C_* and *V_Cm_*, respectively) where *V_Cm_* is also affected by the membrane’s charge dynamics. (**d**) Predicted vs. empirical thresholds (Fig. 5a in ref. 18). The constant a determined from the best fit was used to calculate the surface charge difference in rat cortical neurons (solid blue line, Δσ= σ_i_−σ_o_=-0.107 C/m^2^); predictions for an alternative model (Δ σ=−0.05 C/m^2^, dashed red line) are also shown for comparison. σ_i_≈−0.12 C/m^2^ and σ_o_≈−0.013 C/m^2^ for V_cm_ and V_S.C_ the calculation in *(c).* Error bars are ±s.e.

## Discussion

This study explored the effects of temperature changes on membrane capacitance and its associated currents in a joint attempt to clarify the experimental results of a key recent study^16^ and to pave the way towards predictive modeling of INS^2–15^ and other thermal neurostimulation techniques^18–20^, which could potentially facilitate the development of more advanced and multimodal methods for neural circuit control. Another key motivation to pursue this problem came from our noting the very similar temperature-related capacitance rates of change observed in very different model systems (Fig. 1a) suggesting that this value is putatively universal.

Although the pioneering study by Shapiro *et al*.^16^ attempted to unravel their results’ underlying biophysical mechanism, their explanation raised a crucial question that had to be theoretically reexamined: how is it possible that a rise in membrane capacitance with temperature is caused by ion redistribution in the extra-membranal regions, when the reduction in the medium’s dielectric constant and an increase in diffusion/electrical forces should cause an opposite effect? Indeed, upon further scrutiny we found that the peripheral capacitances (bulk and Stern) *decrease* with temperature, a misinterpretation likely attributed to their naïve misdefinition of the membrane mobile charge sign (see Methods). Our model leads to a diametrically different qualitative story, where the underlying phenomenon is now instead attributed to two completely novel mechanisms:

1. Thermal membranal dimensional changes inferred from small-angle x-ray and neutron scattering measurements, which lead to an increase in the overall capacitance and to capacitive displacement currents [*I_m_ = dC_m_/dt* (*V_m_−V_s.c._*), Fig. 3a,b]. These dimensional changes have been observed in a range of different model membranes, including POPC, SOPC, DPhyPC and more^25–27^ and are putatively attributed to entropicaly-driven shortening of the phospholipids’ tails (Fig. 1b), a process termed trans-gauche isomerization^26^. The predicted associated capacitance rates are relatively insensitive to the specific choice of a membrane (0.33±0.05 %/°C for SOPC and 0.32±0.06 %/°C for DPhyPC) and to the baseline temperature^26^.
2. thermally-mediated displacement currents (*Im = −C_m_dVs.c./dt*) resulting from an increase of the surface-charge related potential (Fig. 3b).

This dual mechanism, found by separately considering the temperature-dependent behavior in each membrane sub-region, explains the experimental shift of membrane current when multivalent ions (Mg^2+^ and Gd^3+^) that affect the surface charge^36^ were added to one side of the membranes’ solutions^16^, while almost no effect was observed on membrane capacitance variations (slope in Fig. 3e,g). It also explains the proportionality constant relating the temperature’s time derivative and induced current (see Methods), allowing to estimate a surface charge difference that was found to be in accord with empirical values for neural cells^35^. Despite its simplifying assumptions, the theoretical analysis was found to qualitatively and quantitatively explain the results of artificial membrane, oocyte and cortical culture experiments, using parameters which were taken ‘as is’ from the literature, whenever attainable (Figs. 3 and 4). The identified effects are dominant in the short timescales considered here, but in conjunction with parallel thermal effects like changes in the Q_10_ factor^37^, probably play a significant role under a wide array of thermal modulation scenarios^38^. Related insights may also help guide our understanding of other emerging neuro-physical modalities like magneto-genetic stimulation (whose biophysics is still poorly understood^39, 40^). For example, we note that membrane mechano-electrical effects involving dimensional changes were suggested in other contexts involving changes in intra-membranal forces, including action potential-related intramembrane thickness variations^41–43^ and ultrasound-induced formation of intramembrane cavities (or ‘bilayer sonophores’^44^). The neuronal intramembrane cavitation excitation (NICE) theoretical framework, putatively explains ultrasonic neuromodulation phenomena (suppression and excitation^45^) and predicts the results of a significant number of related experimental studies^46, 47^.

## Methods

### Constitutive electrostatic laws and equations

The relation between the mobile (*Q*) and fixed (–*σ*) membrane charge densities and the potentials (Δ Φ) that fall on the bulk regions (Fig. 2a) is obtained through electrostatic Boltzmann equations^16, 33^:

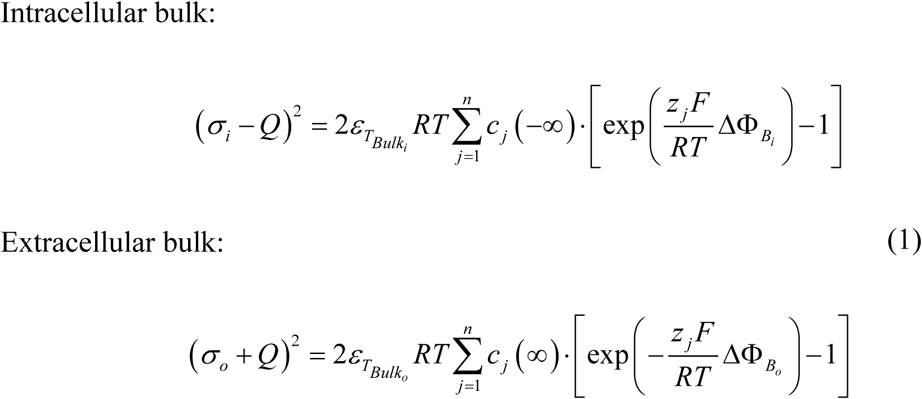

where *ɛ_T_Bulk_i___* and *ɛ_T_Bulk_o___* are the temperature-dependent intracellular and extracellular dielectric constants, respectively; *R* and *F* are the ideal gas and Faraday constants, respectively; *T* is the membrane’s surrounding temperature; *c_j_* (−∞) and *c_j_* (∞) are the intracellular and extracellular ion concentrations far from the membrane; and *z_j_* is ion valence.

The relation between membrane charge and the potentials that fall on the Stern regions can be obtained through the following mathematical formulations^16, 34^, which take into account the respective dielectric constants of the subdivisions of the stern layer (polar heads and ionic region):

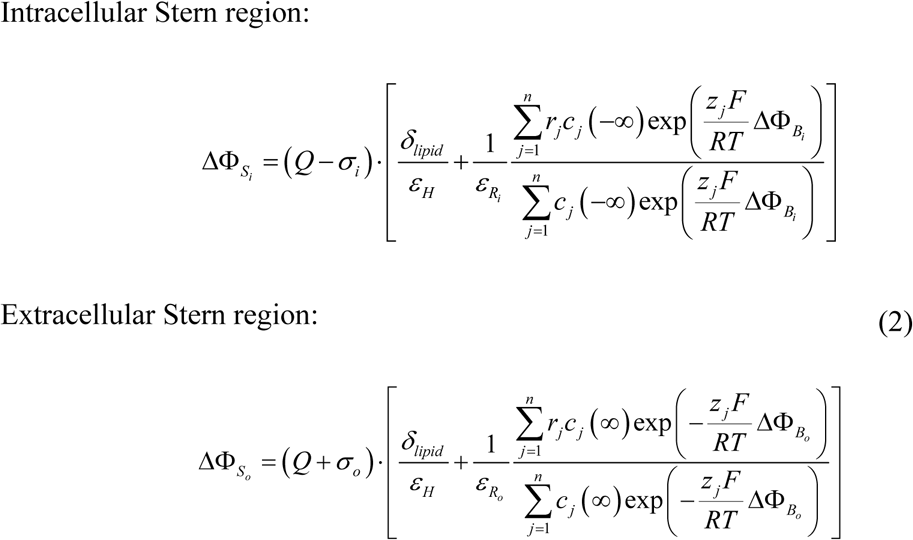

where Δ_Φ*_S_i__*_ and Δ_Φ*_S_o__*_ are the intracellular and extracellular potentials, respectively, that fall on the Stern layers (Fig. 2a); *r_j_* is the ionic effective radius and *ɛ_R_i__*, *ɛ_R_o__* and *ε_H_* are the temperature-dependent dielectric constants in the intracellular and extracellular ionic regions and in the lipid polar heads, respectively. We took *ɛ_R_* = *ɛ_T_Bulk__*, [which is plausible for surface charge density <0.2 C/m^2^ (ref. 48) – the maximal charge density for fluid phase membranes^36^] and 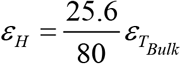 (ref. 49).

The relation between the membrane’s mobile charge and the potential that falls on the hydrophobic core region is linear, under the assumption of zero density of free charges^33^:

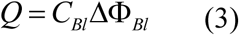

Where *C_Bl_* and ΔΦ_*Bl*_ are the hydrophobic region capacitance and potential. Interestingly, Eqs. 1 and 2 can be seen to yield additional approximately linear relations for the four external membrane regions (see Supplementary Fig. 1), wherein:

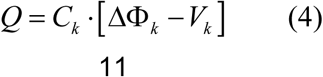

Δ Φ_*k*_ is the potential that falls on each membrane region and *C_k_* and *V_k_* are their respective capacitances and surface charge-related potentials (*k*∈*B_i_*, *S_i_*, *Bl, S_o_* or *B_o_*; 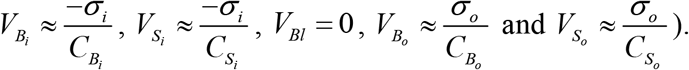

Following the relations shown above the membrane can be represented as a lumped circuit of 5 capacitors with surface charge-related sources in series, where *C_k_* (*T)* and *V_k_* (*T*) are temperature-dependent parameters, which allow us to obtain a simple mathematical linear formula for membrane charge and potential (*V_m_*) (Fig. 2b):

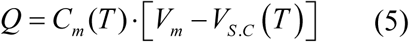

Where 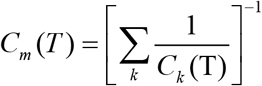 is the total membrane capacitance and 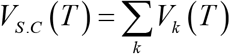. Throughout the study *C_m_*(20 ^*°*^C) was obtained from measurements, and used together with the four capacitance values from Eqs. (1) & (2) to calculate the unknown capacitance *C_Bl_* (except in the theoretical analysis of the temperature effects shown in Figs. 1 and 2, where we used a reference value *C_Bl_* (20°C) calculated in Shapiro *et al.* model).

### Membrane mobile charge notation

Genet *et al*.^33^ defined the extracellular mobile charge as *Q* (*Q*>0) and the intracellular charge as –*Q*. These definitions are opposite to the standard convention, and their naïve application in simulating currents will lead to 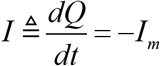 where *I_m_* is a conventional membrane current defined positive for a current flowing from the intracellular to the extracellular domain. It appears that several authors may have made this error^16, 23^. In the results section, the fixed geometry model avoids this notational error (Fig. 2c).

### Dimensional variations

A major conceptual difference between ours and earlier models is the incorporation here of temperature-dependent variations in the membrane’s hydrophobic region dimensions. Estimates of the temperature dependence of the hydrophobic core dimensions were based on published measurements that used X-ray and neutron small-angle scattering experiments^26^, selecting the characteristics of 1-palmitoyl-2-oleoyl-snglycero-3-phosphatidylcholine (POPC), because it relatively closely mimics the mammalian phospholipid composition^50^ (Fig. 1). The hydrophobic core thermal response also modifies Eqs. (1) and (2) by noting that the charge values correspond to areal charge densities.

### Membrane ions absorption

In artificial membrane simulations under Shapiro *et al.* experimental conditions^16^ (symmetrical membranes - 1:1:1 phosphatidylethanolamine (PE):phosphatidylcholine (PC):phosphatidylserine (PS) and 1:1 PE:PC, wherein the PS is a negatively-charged phospholipid molecule, while the PE and PC are nearly uncharged^51^), surface charge change due to ions absorption were modeled through Langmuir isotherm^36^:

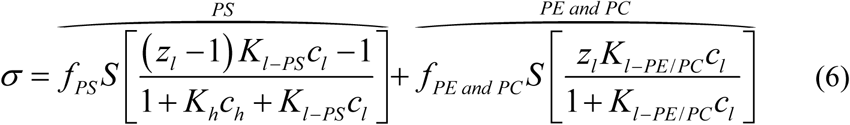

where *S* is the maximal charge density of the ions’ potential membrane binding sites - equals to 0.2 C/m^2^ that corresponds to 0.6nm^2^ per lipid molecule for membranes in fluid phase condition^26, 36^; *f_PS_* and *f_PE and PC_* are the molecular fraction of PS and PE/PC; *z_l_*, *c_l_* and *K_l-PS_* are the valency, concentration close to the membrane and absorption factor to PS of the multivalent *Mg^2+^* or Gd^3+^ ions^16^, respectively; *K_l-PE/PC_* is the multivalent ions’ absorption factor to PE and PC^36, 52^; and *c_h_* and *K_h_* are the concentration and absorption factor to PS of the Na^+^ ions^53^.

### Current and potential calculations

Solving the GCS-based model allowed us to calculate the expected temperature-induced membrane currents in a voltage clamp mode:

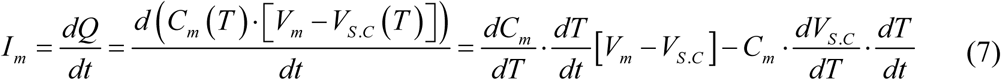

Where 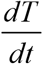 is the temperature time derivative, 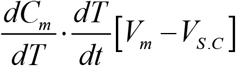 is the membrane capacitive current that arises due to changes in the total membrane capacitance and 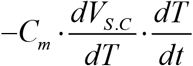 is the surface charge related current.

The membrane potential in a current clamp mode expressed as:

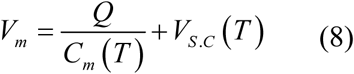

### Estimation of surface charge difference in neurons

To estimate the intracellular and extracellular surface charge difference in neurons, we used our recent discovery that temperature transients excite neurons through formation of membrane currents that proportional to temperature time derivative - 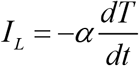, where *α* is known positive constant^18^. This *α* constant can be expressed explicitly by the GCS-based model:

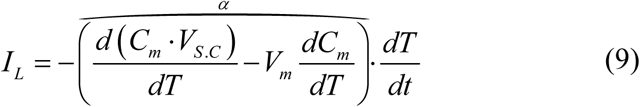

Although *V_m_*dC_m_/dT* is dependent on the membrane potential, its maximal variation of several percent (4 – 7.5 %) prior the action potential formation negligibly affects the *α* value; the main weight of the linear temperature dependent parameter - *C_m_ V_S.C_* (R^2^≥0.999, 20≤T≤50 °C.) explains the emergence of *α* as a constant in Farah *et al.* temperature rate model^18^. Since the membrane surface charge densities determine the *V_S.C_* (see above), it is possible to extract the neural surface charge density difference by knowing the *α*. For accuracy, the calculations account the variation in *V_m_*dC_m_/dT*, by taking the average of the membrane potential prior the action potential formation.

### Model simulation details

The model simulations were conducted in MATLAB, when the solutions were divided into several steps: 1) sub-region parameters were determined explicitly in temperature steps of 0.03 °C (20-50 °C): *C_Bl_* was extracted from the POPC thermal response [Eqs. (3)], while the *C_k_* and *V_k_* in Stern and bulk regions [Eqs. (4)] were determined from linear regression of the potential-charge relations in Eqs. (1) and (2) (see Supplementary Fig. 1); these relations allowed formulating a simple linear membrane capacitor formula [Eq. (5)]. 2) Although *C_m_* and *V_S.C_* demonstrate quite linear temperature-dependence (R^2^≥0.999 and R^2^≥0.998, respectively, 20-50 °C), quadratic curve fittings were implemented for obtaining accurate temperature-dependent analytic formulas. 3) Determination of *C_m_* and *V_S.C_* formulas led to simulations of temperature transient-related membrane currents and potentials [Eqs. (7) and (8)].

IR-induced temperature dynamics was extracted indirectly from Shapiro *et al.* simulations^16^ (laser energy - 7.3 mJ; duration - 10 ms), wherein currents are proportional to temperature time derivative (Fig. 4c of their study):

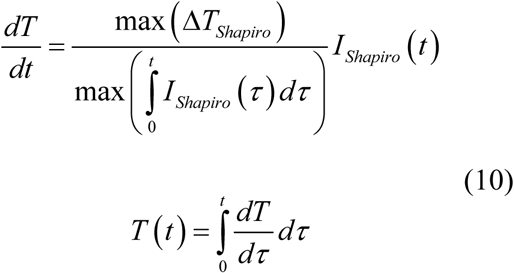

when max (Δ*T_shapiro_*) = 22.2 *°C* is the maximal temperature associated with laser energy - 7.3 mJ and duration - 10 ms.

Finally, the average artificial membrane capacitance in Shapiro *et al.* experiments at a reference temperature of 20°C was determined from charge-potential slopes (Figs. 3c and 4f of their study, obtained by linear regression) and maximal capacitance change percentage (Fig. 3e of their study) for energy - 7.3 mJ and duration - 10 ms; the slopes represent temperature-induced maximal capacitance change [seen from integration of Eq. (7)]. The reference artificial membrane capacitance obtained from this analysis is 89±5.5 pF.

### Statistical analysis

The reported errors in text and figures are s.e.m., unless it is stated otherwise.

## Acknowledgments

This work was supported by an ERC COG #648927 grant, the Russell Berrie Nanotechnology Institute, and an Israel Science Foundation grant No. 1725/13. We thank Shimon Marom, Dmitry Rinberg, Sarah Shaykevich, and Shelly Tzlil for helpful comments.

## Author contributions

M.P and S.S designed the research. M.P performed the simulations and analyzed the data with S.S. S.S and E.K supervised the project and prepared the manuscript with M.P.

